# Batch Action PoTential Analyser (BAPTA): an open source tool for automated high throughput analysis of cardiac action potentials

**DOI:** 10.1101/2023.03.01.530103

**Authors:** Vladislav Leonov, Eleonora Torre, Carlotta Ronchi, Lia Crotti, Peter J Schwartz, Marcella Rocchetti, Antonio Zaza, Luca Sala

## Abstract

The cardiac action potential (AP) is a key species-specific feature of cardiomyocytes that occurs in response to coordinated actions of ion channels. It represents the first step of the cardiac excitation-contraction coupling and it is crucial for cardiomyocyte (CM) physiology. Changes in the cardiac AP may primarily occur as a consequence of diseases or as a direct or unwanted response to drugs. Our ability to quantify these changes defines the reliability of our measurements and its throughput.

Cardiac AP parameters are often quantified through manual time-consuming data analysis protocols or custom-made and proprietary data analysis pipelines; to the best of our knowledge, no tools are currently available for automated cardiac AP analysis and AP parameter quantification.

Here we introduce a free and open source software tool named Batch Action PoTential Analyser (BAPTA), written in the R language, designed to i) overcome the inherent operator-dependent bias on trace selection affecting reproducibility, ii) vastly improve the throughput of the analyses of large datasets and iii) analyse both spontaneous and triggered APs from CMs of multiple species and origin.

We present here four use-cases in which BAPTA can be used at high throughput to investigate the effects of: 1) a disease (cardiomyopathy) on rat CMs, 2) drugs on mouse pacemaker cells, 3) rate-dependency of AP duration in guinea pig CMs and 4) metabolic electrophysiological maturation in human stem-cell-derived CMs. Overall, BAPTA consistently provides faster, more reproducible and scalable readouts which excellently correlate with manual analyses performed by experienced electrophysiologists.

## Introduction

The cardiac action potential (AP) is a species-specific feature of cardiomyocytes (CMs) which originates from the coordinated interplay of ion channels, pumps and ion exchangers. The AP triggers a synchronous calcium release from the sarcoplasmic reticulum and the subsequent contraction, in a process defined excitation-contraction coupling [1]. Deviations from the physiological AP shape profoundly impact the excitation-contraction coupling and may severely affect the heart function. The cardiac AP shape may be altered as results of pathological conditions directly impacting the expression or the biophysical properties of ion channels on the plasma membrane, as occurring in cardiac channelopathies such as the Long QT Syndrome, as a direct consequence of anti-arrhythmic drugs designed to target specific AP phases, or as cardiotoxic or pro-arrhythmic side-effects of drugs [2–4]. The quantitative analysis of cardiac APs is thus pivotal to characterize disease phenotypes and to evaluate drug responses.

Current methods to quantify the main features of cardiac AP rely on manual analyses or proprietary and closed-source software tools, with the consequence of generating costly, operator-dependent and low throughput analyses; this has the significant disadvantage of excluding large portions of academia from accessing these performant analysis tools. Manual analyses, currently used in the vast majority of laboratories, poorly fit with the increasing complexity deriving from high throughput capabilities and, particularly for projects spanning over multiple years, hinder the robustness of research results. Unbiased and automated analyses become also particularly relevant for the most recent and advanced applications generating large amounts of data with high-throughput, high-content or multi-parameter technologies recently emerging to characterize the functional properties of CMs [5,6]. The same strategy has been successful for our previous tool, MUSCLEMOTION [7], in the analysis of contraction profiles from high-speed movies of beating CMs from multiple species. Examples currently exist for the analysis of neuronal APs [8], sinoatrial node cells [9] and optical mapping in whole hearts [10,11], but a solution for atrial and ventricular single cell data in multiple species is currently lacking.

Here, we propose an open source software tool named Batch Action PoTential Analyser (BAPTA), written in the R language[12], which is capable of automatically extracting key features of APs from spontaneously beating or electrically stimulated (paced) CMs from multiple species at high-throughput.

We extensively validated BAPTA with four different use-cases to assess: 1) pharmacological effects on spontaneously-beating sinoatrial node (SAN) CMs derived from mice; 2) electrophysiological changes on AP caused by a cardiomyopathy in rat adult ventricular CMs; 3) the rate-dependence of AP properties in adult guinea pig CMs; 4) the maturation status of ventricular-like human induced pluripotent stem cell-derived cardiomyocytes (hiPSC-CMs).

Across 4 different laboratories, we have used primary CMs from animal species characterized by remarkably different AP shapes (Figure 1), demonstrating a very broad range of AP profiles and applications that cover the majority of applications in the cardiac field for both academia and industry.

**Figure 1:**
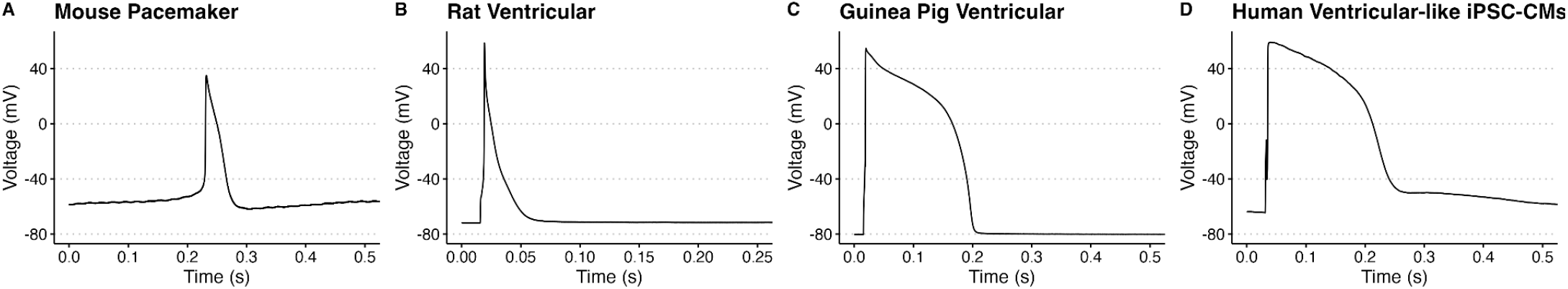
Cardiac APs from A) mouse sinoatrial node cells, B) rat left ventricular CMs, C) guinea pig left ventricular CMs and D) ventricular-like hiPSC-CMs.

Compared to manual analyses performed by experienced electrophysiologists, BAPTA provides rapid, unbiased and reproducible readouts of cardiac APs at high throughput, significantly reducing the time required for the analysis while maintaining high correlations in all the AP parameters considered in the study. Being open source, BAPTA is a dynamic platform that can be expanded, improved, and embedded for customized applications.

## Methods

### Code

BAPTA source code is open source and has been written in the R Language [12]. The user interface has been built with the Shiny package (https://shiny.rstudio.com/). The source code and a detailed user manual are available for use and further development in our GitHub repository (https://github.com/l-sala/BAPTA). The blueprint of BAPTA is visible in Figure 2.

**Figure 2:**
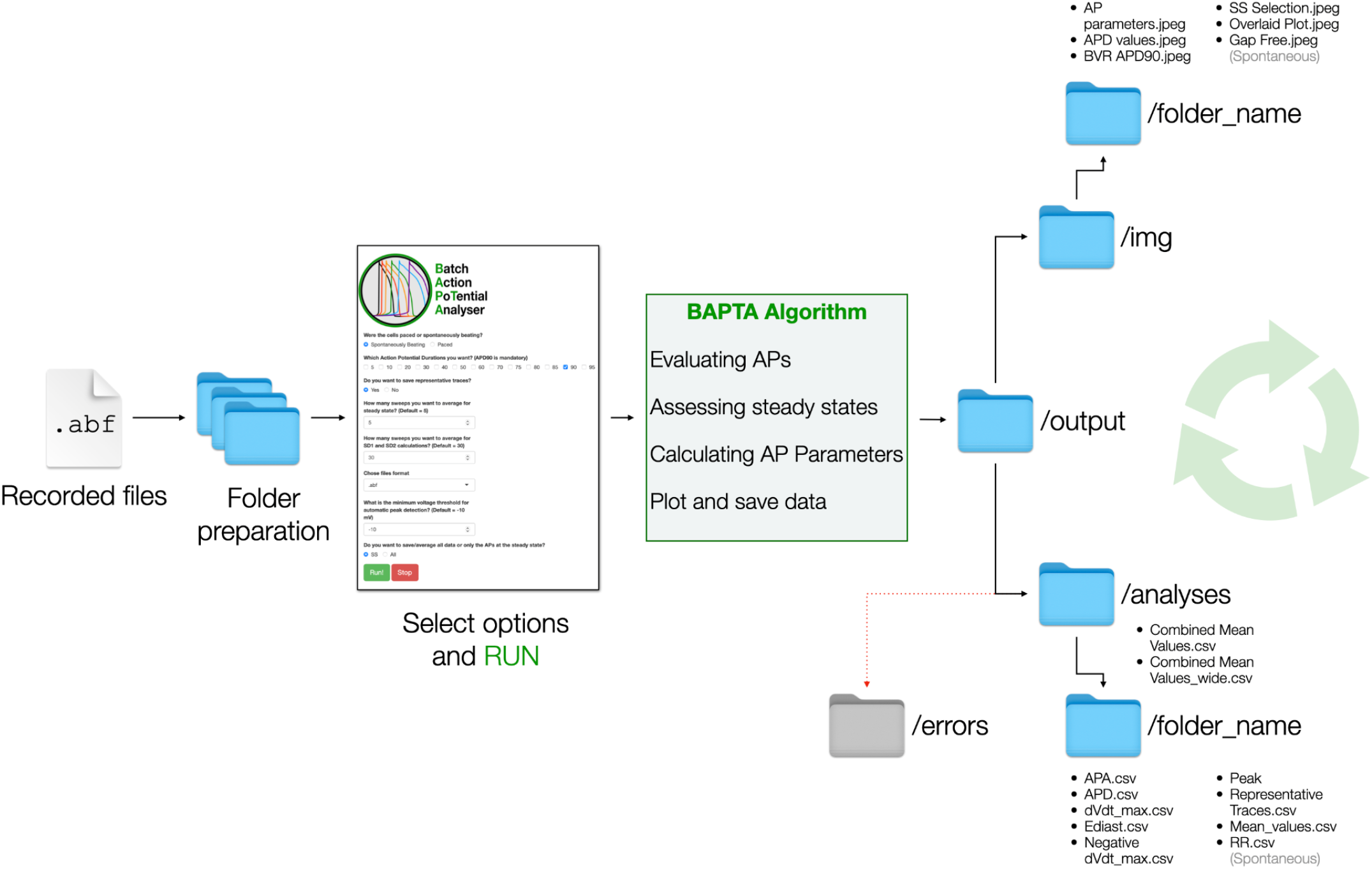
Schematic representation of BAPTA. Abf files recorded with patch clamp are first placed into dedicated folders. Then, options for AP analysis are selected, the files analyzed by BAPTA and the outputs are saved as images or tables (.csv) in the appropriate folders.

### Ethics

All the experiments were conducted according to the Helsinki declaration. The use of animal tissue has been authorized by the Ethical Committee of the University of Milano - Bicocca, through protocols N. 29C09.26 and 29C09.N.YRR approved in January 2021 and June 2018 for rats and guinea pigs respectively and by the Ethical Committee of the University of Montpellier and the French Ministry of Agriculture with protocol N. 2017010310594939.

### Isolation of mouse pacemaker cells

SAN pacemaker cells were isolated from male WT mice hearts as previously described[13]. Briefly, the heart was excised and immersed in a pre-warmed (36 °C) Tyrode’s solution containing 140 mM NaCl, 5.4 mM KCl, 1 mM MgCl_2_, 1.8 mM CaCl_2_, 5.5 mM D-glucose, and 5 mM HEPES (adjusted to pH 7.4 with NaOH). The SAN region was identified using the superior and inferior vena cava, the crista terminalis, and the interatrial septum as landmarks. SAN tissue was excised and transferred into a low-Ca^2+^ solution containing 140 mM NaCl, 5.4 mM KCl, 0.5 mM MgCl_2_, 0.2 mM CaCl_2_, 1.2 mM KH_2_PO_4_, 50 mM taurine, 5.5 mM D-glucose, 1 mg/mL BSA, and 5 mM Hepes–NaOH (adjusted to pH 6.9 with NaOH) for 4 min. Then, enzymatic digestion was carried out for 20-25 min at 36 °C in the low-Ca^2+^ solution containing Liberase TH Research Grade (0.15 mg/mL; Roche) and elastase (0.5 mg/mL; Worthington). Digestion was stopped by washing the SAN in a “Kraftbrühe” (KB) medium containing 100 mM K-glutamate, 10 mM K-aspartate, 25 mM KCl, 10 mM KH_2_PO_4_, 2 mM MgSO_4_, 20 mM taurine, 5 mM creatine, 0.5 mM EGTA, 20 mM D-glucose, 5 mM HEPES, and 1 mg/mL BSA (adjusted to pH 7.2 with KOH). Single SAN cells were then dissociated from the SAN tissue by manual agitation using a flame-forged Pasteur’s pipette. To recover the automaticity of the SAN cells, Ca^2+^ was gradually reintroduced in the cell’s storage solution to a final concentration of 1.8 mM. SAN cells were then left to rest for 1 h before recordings.

### Type 1 diabetes (T1D) cardiomyopathy rat model

Male Sprague Dawley rats (150–175 gr) were used to generate a streptozotocin (STZ)-induced T1D cardiomyopathy model. T1D was induced through a single STZ (50 mg/Kg) injection into a rat-tail vein; littermate control (healthy) rats received only citrate buffer (vehicle). STZ-treated (diseased) rats showed diastolic dysfunction (DD) as previously published[14].

### Isolation of rat and guinea pig CMs

Adult healthy and diseased rats and guinea pig ventricular CMs were isolated as previously described [15][16]. Briefly, the heart was excised and immersed in a pre-warmed (36 °C) “base” solution containing 143 mM NaCl, 5.4 mM KCl, 0.5 mM MgCl_2_, 1.8 mM CaCl_2_, 0.25 mM NaH_2_PO_4_, 5.5 mM D-glucose and 5 mM Hepes–NaOH (adjusted to pH 7.4 with NaOH). By using a Langendorff column for retrograde coronary perfusion, the heart was perfused with a warm (36 °C) “base” solution. This perfusion was maintained until vigorous mechanical activity resumed and blood and clots were completely removed. The heart was then perfused, until arrest occurred, with a Ca^2+^-free “base” solution. Then, enzymatic digestion was carried out for 3-5 min for rats and 1-2 min for GPs at 36 °C perfusing the heart with “Kraftbrühe” (KB) medium containing 70 mM KOH, 50 mM glutamic-acid, 40 mM KCl, 20 mM KH_2_PO_4_, 3 mM MgCl_2_, 20 mM taurine, 0.5 mM EGTA-KOH, 10 mM D-glucose and 10 mM HEPES (adjusted to pH 7.4 with KOH). Liberase TM Research Grade (0.1 mg/mL; Roche, Germany), Trypsin-EDTA 1x (Euroclone, Italy) and 6.5 µM CaCl_2_ were added to KB medium to digest the heart. Digestion was stopped by washing the heart with enzymes-free KB medium. The atria were dissected, and the ventricles were chopped to 1-mm fragments. The fragments were exposed to gentle mechanical agitation in KB medium. Samples of the suspension were collected, filtered through a 75µm-width nylon mesh, and centrifuged at 500 rpm for 3 minutes. The pellets were resuspended in KB medium and stored in this solution at 4 °C. Before recordings, Ca^2+^ was gradually reintroduced in the cell’s storage solution to a final concentration of 0.8 mM. Rod-shaped, Ca^2+^ -tolerant myocytes were used within 12 h from dissociation.

### Differentiation of hiPSCs to hiPSC-CMs

The following hiPSC line was obtained from the NIGMS Human Genetic Cell Repository at the Coriell Institute for Medical Research: GM25256. hiPSCs were cultured on recombinant human vitronectin (rhVTN, ThermoFisher) in E8 Flex medium (ThermoFisher) and differentiated on cell culture-grade Matrigel (BD). Ventricular-like hiPSC-CMs were differentiated from hiPSCs by using a previously published protocol based on the modulation of the Wnt-signalling pathway [17], purified through glucose starvation (> 90% CMs) and cryopreserved in serum-free 10% DMSO-based cryopreservation medium before day 20. Cryopreserved hiPSC-CMs were thawed before each experiment and maintained in culture for two weeks in RPMI medium supplemented with B27 Supplement. The metabolic maturation medium was a RPMI1640-derivation supplemented with fatty acid according to a previously published protocol [18].

### Electrophysiology

SAN cells were seeded on custom-made chambers with 20 mm Ø glass bottoms for cell attachment with a density of 50-100 cells/cm^2^ and patched with a Molecular Devices 700A amplifier coupled with a Molecular Devices 1550B Digidata.

The extracellular solution (Tyrode’s based) contained (mM): 140 NaCl, 5.4 KCl, 1 MgCl_2_, 1.8 CaCl_2_, 5 HEPES, 5.5 D-Glucose; pH was set to 7.4 with NaOH.

Glass pipettes were filled with the intracellular solution contained (mM): 80 K-Aspartate, 50 KCl, 1 MgCl_2_, 5 HEPES, 2 CaCl_2_, 5 EGTA, 3 ATP-Na^+^ salt; pH was adjusted to 7.2 with KOH. The intracellular solution was cryopreserved at -20 °C and thawed before each experiment. Escin was dissolved in sterile H_2_O and added to the intracellular solution (40 µg/mL) right before each experiment.

Measurements were performed in perfused cells at physiological temperature (37 °C) with a temperature-controlled chamber and warmed solutions.

Rat and GP ventricular CMs were seeded on 35 mm Ø culture dishes with a density of 500 cells/cm^2^ and patched with a Molecular Devices 200A amplifier coupled with a Molecular Devices 1200 Axon Digidata.

The extracellular solution (Tyrode’s based) contained (mM): 154 NaCl, 4 KCl, 1 MgCl_2_, 2 CaCl_2_, 5 HEPES-NaOH, 5.5 D-Glucose; pH was set to 7.35 with NaOH.

Glass pipettes were filled with the intracellular solution contained (mM): 110 K-Aspartate, 23 KCl, 3 MgCl_2_, 5 HEPES-KOH, 0.2 CaCl_2_, 0.5 EGTA-KOH, 0.4 mM GTP-Na^+^ salt, 5 mM ATP-Na^+^ salt, and 5 mM creatine phosphate Na^+^ salt; pH was adjusted to 7.3 with KOH. The intracellular solution was cryopreserved at -20 °C and thawed before each experiment.

hiPSC-CMs were seeded on matrigel-coated 10 mm Ø glass coverslips with a density of 8.5 - 12.5 * 10^3^ cells/cm^2^ and patched with a Molecular Devices 200B amplifier coupled with a Molecular Devices 1440A Digidata.

The extracellular solution (Tyrode’s based) contained (mM): 154 NaCl, 5.4 KCl, 1 MgCl2, 1.8 CaCl_2_, 5 HEPES, 5.5 D-Glucose; pH was set to 7.35 with NaOH.

Glass pipettes were filled with the intracellular solution contained (mM): 125 K-Gluconate, 20 KCl, 10 NaCl, 10 HEPES; pH was adjusted to 7.2 with KOH. The intracellular solution was cryopreserved at -20 °C and thawed before each experiment. Amphotericin B was dissolved in DMSO to a final concentration of 0.22 mM and added to the intracellular solution right before each experiment.

Measurements were performed in perfused cells at physiological temperature (37 °C) with a temperature-controlled chamber and warmed solutions.

Manual analyses were performed by experienced electrophysiologists with Clampfit (Molecular Devices) to obtain key parameters of the cardiac action potential described in Table 1.

**Table 1.**
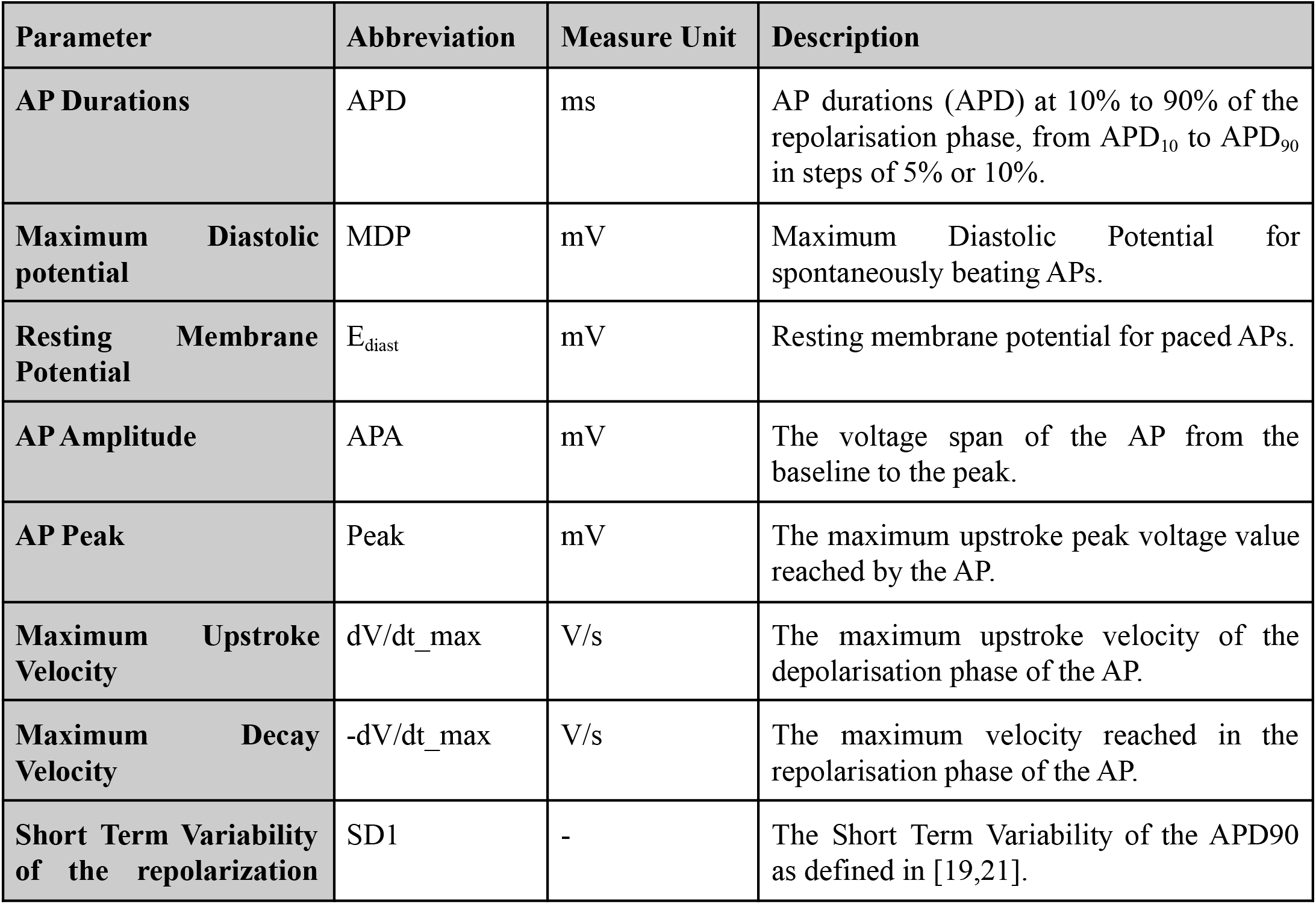
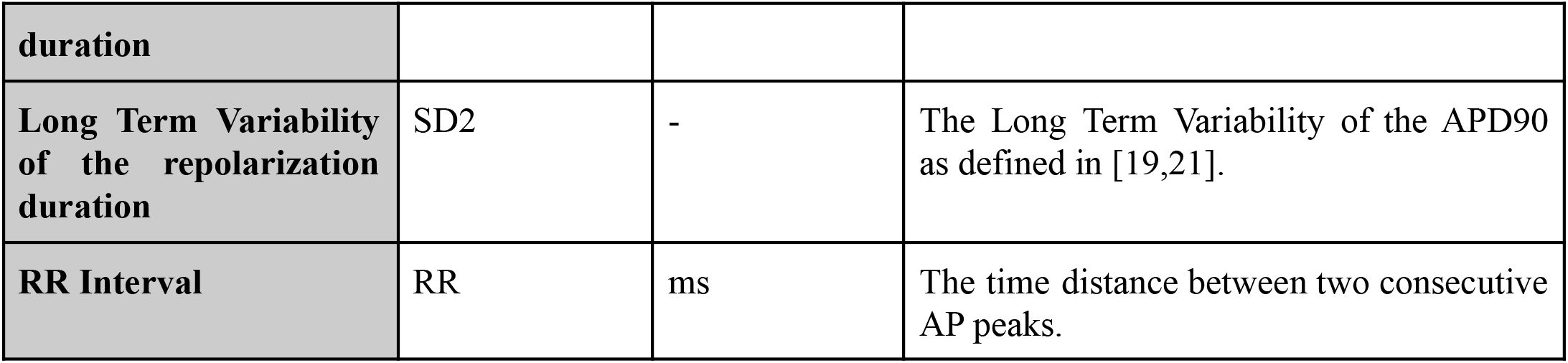
Action Potential parameters outputed by the software

| Parameter | Abbreviation | Measure Unit | Description |
| --- | --- | --- | --- |
| AP Durations | APD | ms | AP durations (APD) at 10% to 90% of the repolarisation phase, from $APD_{10}$ to $APD_{90}$ in steps of 5% or 10%. |
| Maximum Diastolic potential | MDP | mV | Maximum Diastolic Potential for spontaneously beating APs. |
| Resting Membrane Potential | $E_{diast}$ | mV | Resting membrane potential for paced APs. |
| AP Amplitude | APA | mV | The voltage span of the AP from the baseline to the peak. |
| AP Peak | Peak | mV | The maximum upstroke peak voltage value reached by the AP. |
| Maximum Upstroke Velocity | $dV/dt_{max}$ | V/s | The maximum upstroke velocity of the depolarisation phase of the AP. |
| Maximum Decay Velocity | $-dV/dt_{max}$ | V/s | The maximum velocity reached in the repolarisation phase of the AP. |
| Short Term Variability of the repolarization | SD1 | - | The Short Term Variability of the $APD_{90}$ as defined in [19,21]. |
| **duration** |  |  |  |
| **Long Term Variability of the repolarization duration** | SD2 | - | The Long Term Variability of the APD90 as defined in [19,21]. |
| **RR Interval** | RR | ms | The time distance between two consecutive AP peaks. |

### Steady State Selection

The identification of a steady state is crucial in each experiment to increase the accuracy and reproducibility of the measurements. A steady state is reached when a sufficient number of consecutive beats exhibits consistent and stable values; the steady state and the number of beats in it is often operator-dependent and is evaluated live during the recording. The steady-state is further manually re-assessed by the operator during the analyses and a fixed number of beats will be averaged to obtain mean values for each parameter. BAPTA fully automates this process with different strategies based on the experiment type (paced or spontaneous).

#### Paced CMs

In paced CMs, the steady state was selected according to two different criteria, which were prioritized as follows: 1) Stability of APD_90_ among N consecutive beats; 2) E_diast_ polarization among the N APs with the most hyperpolarized values. The N of APs to be selected is customizable by the user in the interface. Among the groups of N APs with the lowest APD_90_ variability, BAPTA selects those which have the most polarized (i.e. negative) E_diast_. This excludes a significant number of artifacts and guarantees the selection of properly-polarized APs in case of challenging recordings with unstable APDs or E_diast_. The selection of properly polarized APs also minimizes inclusion of experimental artifacts in the analyses, such as those that arise as a consequence of a lost seal. The same function, on a customizable number of beats, is used to select a longer interval for the calculation of the orthogonal- (SD1) and longitudinal (SD2) standard deviation of the Poincaré plot of APDs [19].

#### Spontaneously beating CMs

In spontaneously beating CMs, the identification of a Maximum Diastolic Potential (MDP) and APDs are more challenging, particularly in situations where Vm oscillations significantly alter the AP contour. The automatic detection of peaks was achieved through iterative steps aimed to improve the accuracy of the readout. BAPTA identifies peaks with the *findpeaks* function in the *pracma* package [20], using a threshold minimum peak height which can be customized by the user in the interface to exclude -if required-every non-physiological alteration of the membrane potential that cannot be attributed to the AP. An additional level is introduced to ensure that all the peaks are picked up correctly by looping through all the values above the minimum peak height to verify whether they are the true peaks or artifacts. BAPTA also filters every value above +70 mV to remove unphysiological electrical artifacts during peak identification. Once the coordinates of peaks have been identified, the automatic detection iterates to define the most negative Vm values among them as MDP. Outliers such as voltage noise spikes or rapid Vm changes occurring from loose seals or detached cells (i.e. voltage drops to 0) within a file are excluded by selecting physiological ranges of voltage for cardiac AP. Once the spontaneous APs have been separated and sorted, they are processed as for paced CMs.

### Determination of AP parameters

For paced APs, the peaks (Peak) are defined as the maximum voltage values after the passive membrane potential change induced by the stimulation current occurring in the first 10% of the AP. In case of spontaneously beating CMs, peaks are identified through a dedicated function as described above. E_diast_ was defined as the mean of the first 10 points occurring before the AP stimulation artifact in case of paced APs while they were defined as the minimum point before the AP in case of spontaneous APs. AP Amplitude (APA) is defined as the difference between the Peak voltage value and the E_diast_ voltage value. Positive (dV/dt_max) and negative (-dV/dt_max) upstroke velocities were defined as the coordinates at which the fastest depolarization and repolarization occur; they are respectively calculated as the first derivative of the AP upstroke and the first derivative of the late AP repolarization phase (phase 3 → phase 4). Short- and Long-term variability of the repolarization duration (here respectively termed SD1 and SD2) are important descriptors of APD variability and describe the temporal patterns of APD variation through its short- and long-term components [21]. BAPTA calculates SD1 and SD2 as previously described [19,21] on a fixed number of beats (default = 30); the number of beats can be customized in the user interface settings. Beating frequency is calculated as the mean rate at which spontaneous APs occur as well as the time interval between consecutive peaks, defined RR interval as a reference to the respective clinical parameter in the electrocardiogram.

## Results

### Application of BAPTA to quantify parameters from spontaneously beating mouse pacemaker cells

SAN CMs are characterized by a spontaneous Ca^2+^-driven rhythmic electrical activity, with the activity of funny current (I_f_) and the Ca^2+^-clock as two main drivers of automaticity [22]. It is then important to evaluate the spontaneous beating frequency as well as the AP parameters to understand whether drugs positively or negatively impacting the activity of SAN cells (e.g. Ivabradine, β-adrenergic stimulation)[23,24] or disease conditions (e.g. sick sinus syndrome[25,26]) may affect APs, their relative frequency or temporal variability.

This use case analyses recordings from spontaneously beating CMs, and specifically from mouse SAN CMs, with the aim to extract relevant temporal parameters from spontaneous APs. BAPTA efficiently analysed continuous (*gap free)* recordings of APs from mouse pacemaker cells. The AP temporal parameters obtained by BAPTA well correlated with those analysed manually (Figure 3 A-B), with R^2^ spanning from 0.94 to 0.99. SAN cells were challenged with isoproterenol, which induced positive inotropic and chronotropic effects as expected from beta-adrenergic agonists (Figure 3 C-D); these AP changes were correctly captured by BAPTA.

**Figure 3:**
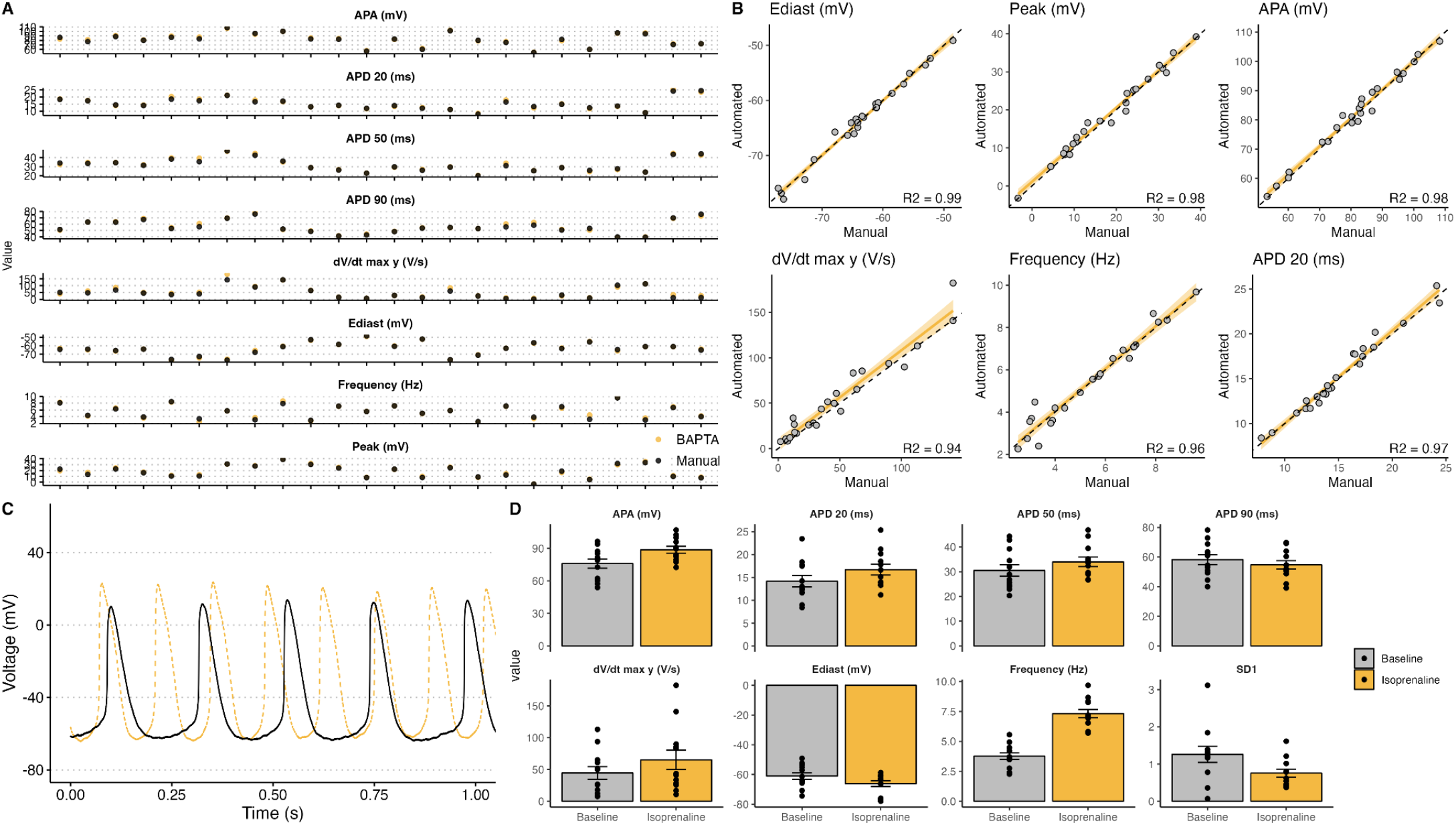
A) Raw values of AP parameters; each dot represents a single recording, with parameters obtained from Manual (black) or Automated (orange) analysis. B) Linear regression of results obtained from manual analyses (x axis) and those obtained with BAPTA (y axis). C) Representative traces from mouse sinoatrial node cardiomyocytes in normal conditions (black) and after superfusion with isoprenaline (orange). D) Mean parameters and individual data points obtained with BAPTA from sinoatrial node cardiomyocytes at baseline (gray) and after superfusion with isoprenaline (orange).

### Application of BAPTA to distinguish healthy vs disease phenotype using adult rat ventricular myocytes

Ventricular APs from rats are characterized by a sharp AP phase 0, a very steep repolarization phase mainly driven by I_Kur_ and I_to_ [27] and a negligible AP plateau. These models are widely used in disease-modelling, toxicology and pharmacology due to their versatility, decent translational relevance and low costs. This use-case compares AP values from ventricular CMs, derived from healthy or diseased rats, with the aim to detect disease-induced changes in the AP shape or parameters. BAPTA successfully detected changes in APDs, APA, dV/dt_max, SD1 induced by the disease. Once correlated with the results obtained from time-consuming manual analyses by an experienced electrophysiologists, we identified excellent correlations for all the parameters with particularly remarkable R^2^ for temporal parameters (APDs, (0.98-0.99) and voltage values (E_diast_, APA, 0.91-0.94) (Figure 4 A-B). APs from diseased rats exhibited markedly prolonged APD_20_, APD_50_ and APD_90_, reduced APA and dV/dt_max and, as expected, an enhanced SD1 (Figure 4 C-D).

**Figure 4:**
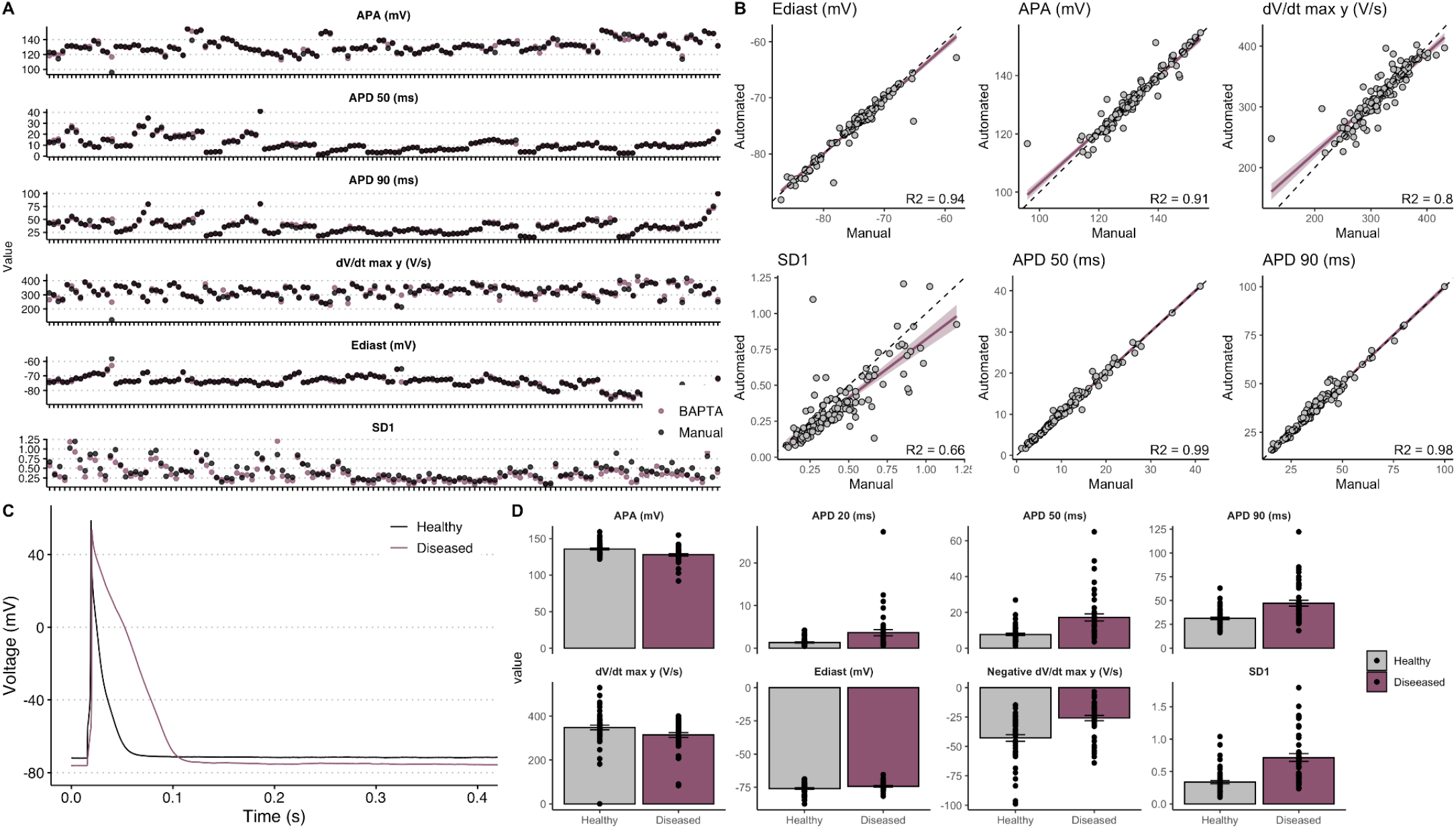
Analysis of disease phenotypes: A) Raw values of AP parameters; each dot represents a single recording, with parameters obtained from Manual (black) or Automated (purple) analysis. B) Linear regression analysis between results obtained from manual analyses (x axis) and those obtained with BAPTA (y axis). C) Representative example of rat ventricular AP from healthy (black) or diseased (purple) rats. D) Mean parameters obtained by BAPTA from healthy (grey) or diseased (purple) rats.

### Application of BAPTA to assess the rate-dependence of APD using adult guinea pig ventricular myocytes

APs from GP ventricular CMs are characterized by a marked plateau phase sustained by I_CaL_ and a repolarization phase driven by I_Kr_ and I_Ks_, with the latter particularly relevant at high pacing rates or under β-adrenergic activation [16,28,29]. GP ventricular CMs represent an optimal tradeoff between costs and translational relevance: APD values are closer to those of human ventricular CMs [30], particularly if compared to those of mice and rats, and have significantly more affordable costs compared to rabbits or dogs.

APs from GPs were recorded with patch clamp in isolated ventricular GP CMs paced at four different frequencies (0.5 Hz, 1 Hz, 2 Hz, 4 Hz) and analysed either manually, with pClamp, or with BAPTA. Representative traces and correlations between manual and automated analyses exhibited very high correlation for all the parameters involved (Figure 5 A-B). The automatic analysis of the rate-dependence of APD exhibited the expected reverse relationship (i.e. the higher the pacing rate, the shorter the APD [28]) (Figure 5 C-D).

**Figure 5:**
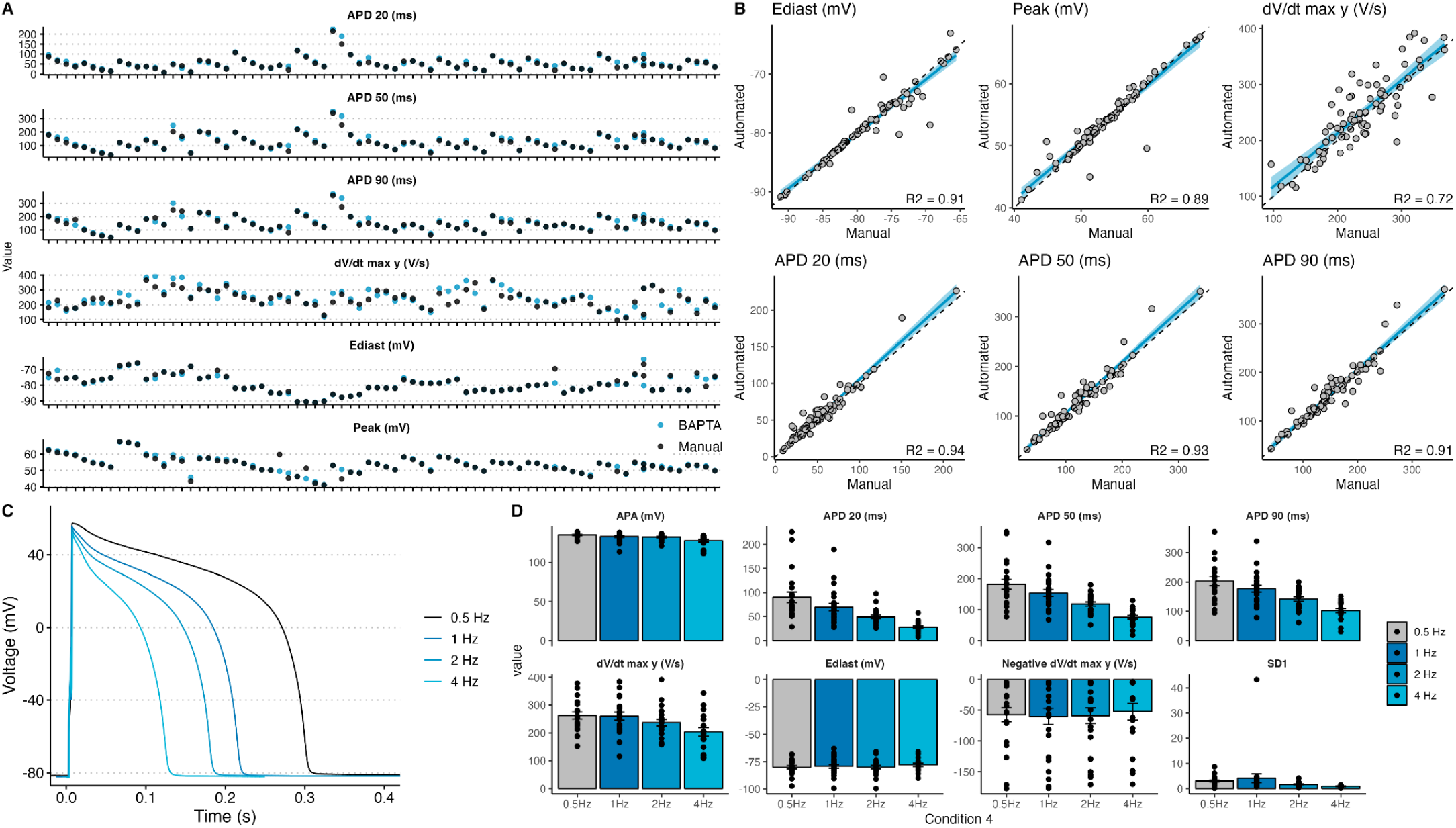
Analysis of rate dependence of APD: A) Raw values of AP parameters obtained from Manual (black) or Automated (blue) analysis. B) Linear regression analysis between results obtained from manual analyses (x axis) and those obtained with BAPTA (y axis). C) Representative examples from guinea pig ventricular CMs stimulated at different pacing frequencies. D) Mean parameters obtained by BAPTA at different pacing rates.

### Application of BAPTA to assess the maturation status of stem cell-derived CMs using human ventricular-like iPSC-CMs

Ventricular-like hiPSC-CMs have a more immature phenotype compared to their adult counterpart when isolated from bidimensional monolayers [30,31]. Multiple strategies to improve hiPSC-CM maturation have been proposed including culture in 3D, co-culture with other relevant cell types [32,33], metabolic maturation [18,34] or electrical stimulation [35,36]. The field strives to identify molecules, cocktails or stimuli to improve hiPSC-CM maturation, and electrophysiological and functional readouts represent mandatory and critical steps to evaluate the efficiency and robustness of these maturation protocols. APs from healthy ventricular-like hiPSC-CMs were recorded at different pacing frequencies and analyzed either manually, with pClamp, or with BAPTA.

Automatic analyses are very well correlated with manual analyses performed by experienced electrophysiologists, with excellent correlations for the 6 critical parameters considered in our analyses. Coefficients of determination for these correlations span from 0.96 to 0.99 (Figure 6 A-B).

**Figure 6:**
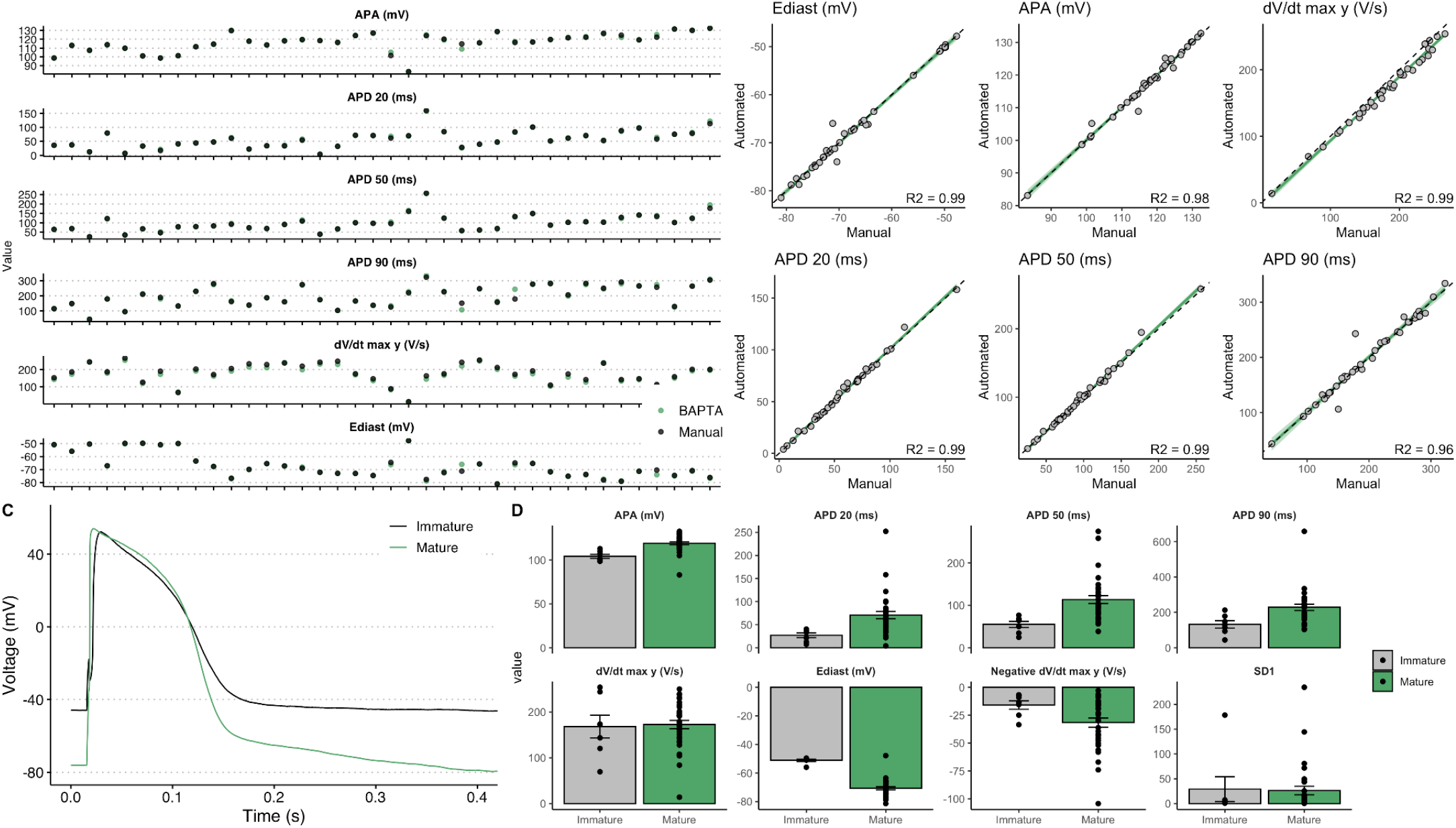
Analysis of hiPSC-CM electrophysiological maturation: A) Raw values of AP parameters obtained from Manual (black) or Automated (green) analysis. B) Linear regression analysis between results obtained from manual analyses (x axis) and those obtained with BAPTA (y axis). C) Representative examples from immature (black) or mature (green) ventricular-like hiPSC-CMs. D) Mean parameters obtained by BAPTA in immature (grey) or mature (green) hiPSC-CMs.

BAPTA efficiently identified that the metabolic maturation protocol used here [18] effectively pushed many electrophysiological parameters from immature values to resembling those of adult CMs. E_diast_ was significantly hyperpolarized by the maturation treatment and, as a consequence, both APA and dV/dt max were improved. APD_20_, APD_50_ and APD_90_ were significantly prolonged, particularly due to the larger recruitment of ion channels in response to the more hyperpolarized E_diast_, while the negative dV/dt max, indicating the steepness of the repolarization phase, had significantly more negative velocity values (i.e. steeper repolarization). No changes were observed in SD1 (Figure 6 C-D).

## Discussion

Efficient and scalable methods to analyse electrophysiological parameters represent a significant benefit for scientists and laboratories working on cardiovascular electrophysiology. Here we have presented BAPTA, an easy-to-use open source software tool for the automatic analysis and quantification of relevant parameters of cardiac APs acquired with patch clamp. Overall, its integration into current research practices will significantly facilitate the assessment of cardiac cellular electrophysiology and will improve data sharing, reproducibility and comparisons of those working in stem cell biology departments lacking expertise in electrophysiology or laboratories aiming to optimize data analysis. The applications for BAPTA are very broad and include: i) the assessment of the effects of drugs on the cardiac APD[3]; ii) the identification of a disease phenotype associated with cardiac channelopathies and in particular with APD shortening, as in the Short QT Syndrome[37], APD prolongation as in the Long QT Syndrome[38], or with changes in the AP contour such as in the Brugada Syndrome[39]; iii) the comparison of species-specific AP features and iv) to rapidly establish the effects of genetic variants of unknown significance[40]. Similarly to the previous software tool MUSCLEMOTION[7], BAPTA can be expanded and integrated into large data analyses pipelines and coupled with other software tools to automatically quantify biological parameters as calcium transients or contractility.

Here, BAPTA was effective in the analysis of APs from the most widely used species and platforms in both academia and industry. The reliability of BAPTA was assessed on a broad range of experimental conditions and on a wide range of AP shapes, making us confident that such capabilities will extend to APs from species not specifically tested here (e.g. APs from zebrafish CMs, adult mouse CMs, human atrial CMs, etc.).

Currently, to the best of our knowledge, open source solutions for the analysis of spontaneous and triggered cardiac APs exist but they are often limited by command-line input interfaces; a few commercial solutions might partially circumvent this limitation, but usually require programming skills, extensive timing for manual data analysis, costly licences and rely on proprietary and closed source software environments. The reliability of BAPTA was assessed against manual analyses from experienced electrophysiologists, obtaining excellent correlations; lower R^2^ were instead obtained for SD1 values, but the reason underlying this discrepancy is intrinsic in the steady-state selection function of BAPTA compared to the method used for manual analyses. BAPTA automatically and reproducibly selects the most stable set of beats on which the SD1 is calculated, while a human operator usually selects those close to the end of the recording. Indeed, manual analyses do not rely on clear, specific and reproducible criteria and suffer from operator-dependent bias. Hence, the lower R^2^ values obtained for SD1 correlations have to be positively interpreted. In all experimental conditions, the automatically calculated dV/dt max appeared slightly overestimated compared to the manual one. The reason behind this is the automatic steady-state selection that prioritizes hyperpolarized APs in stable regions of the recordings rather than using the last sweeps of the recordings as generally performed manually. As previously mentioned for SD1, this systematic behavior is reproducible across different datasets and does not affect the final results in case all datasets were analyzed with the reproducible criteria included in BAPTA.

A massive gain in the time spent for the analyses was also achieved: the manual analyses of the data included in this manuscript took several weeks to be completed, while the very same data could be analysed by BAPTA overnight.

Overall, BAPTA provides quantitative information on relevant cardiac AP parameters in multiple conditions, significantly improves the reproducibility and throughput of current-clamp data analysis and offers solid and efficient operator-independent readouts that perfectly couples with large high throughput screening capabilities.

## Limitations

BAPTA cannot discriminate between normal or diseased action potentials, thus arrhythmic events occurring during phases 1-3 of the cardiac AP (e.g. EADs) are not currently recognised.

BAPTA can currently be used downstream to patch clamp recordings obtained from Molecular Devices hardware and the respective .*abf* files. Files recorded with other systems must be first converted to comma separated value or text files (see manual) before their analysis with BAPTA.

## Data Availability

The source code and the data included in this article are available for use and further development at the following GitHub repository (https://github.com/l-sala/BAPTA). A step-by-step manual is also available on the repository.

## Author Contributions

VL: Wrote and bug-fixed the code, performed experiments in hiPSC-CMs and data analysis.

ET: Performed experiments and analyses in rat CMs and mouse SAN cells.

CR: Performed experiments and analyses in GP CMs.

LC: Provided constructive feedback on the manuscript.

PJS: Provided constructive feedback on the manuscript.

MR: Provided constructive feedback on algorithms, analyses and manuscript. Supervised experiments on Rat CMs.

AZ: Provided constructive feedback on algorithms, analyses and manuscript. Supervised experiments on GP CMs.

LS: Conceptualized the software tool, wrote and bug-fixed the code, supervised experiments in hiPSC-CMs and performed data analysis.

## Supplementary Figures

**Figure S1:**
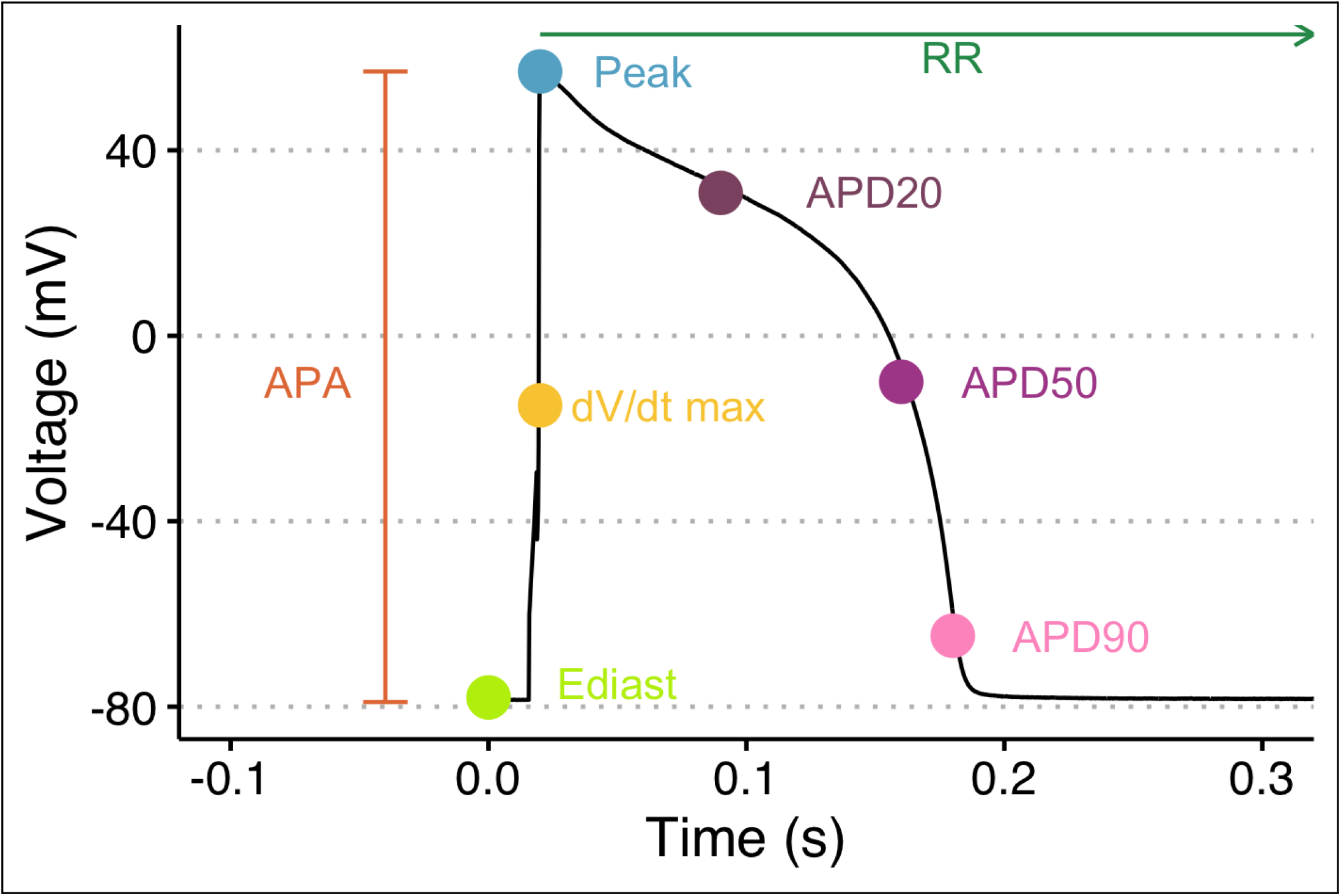
AP parameters automatically extracted by BAPTA.

