## Supplementary figures and images for "Batch Action PoTential Analyser (BAPTA): an open source tool for automated high throughput analysis of cardiac action potentials"

### Figure S1

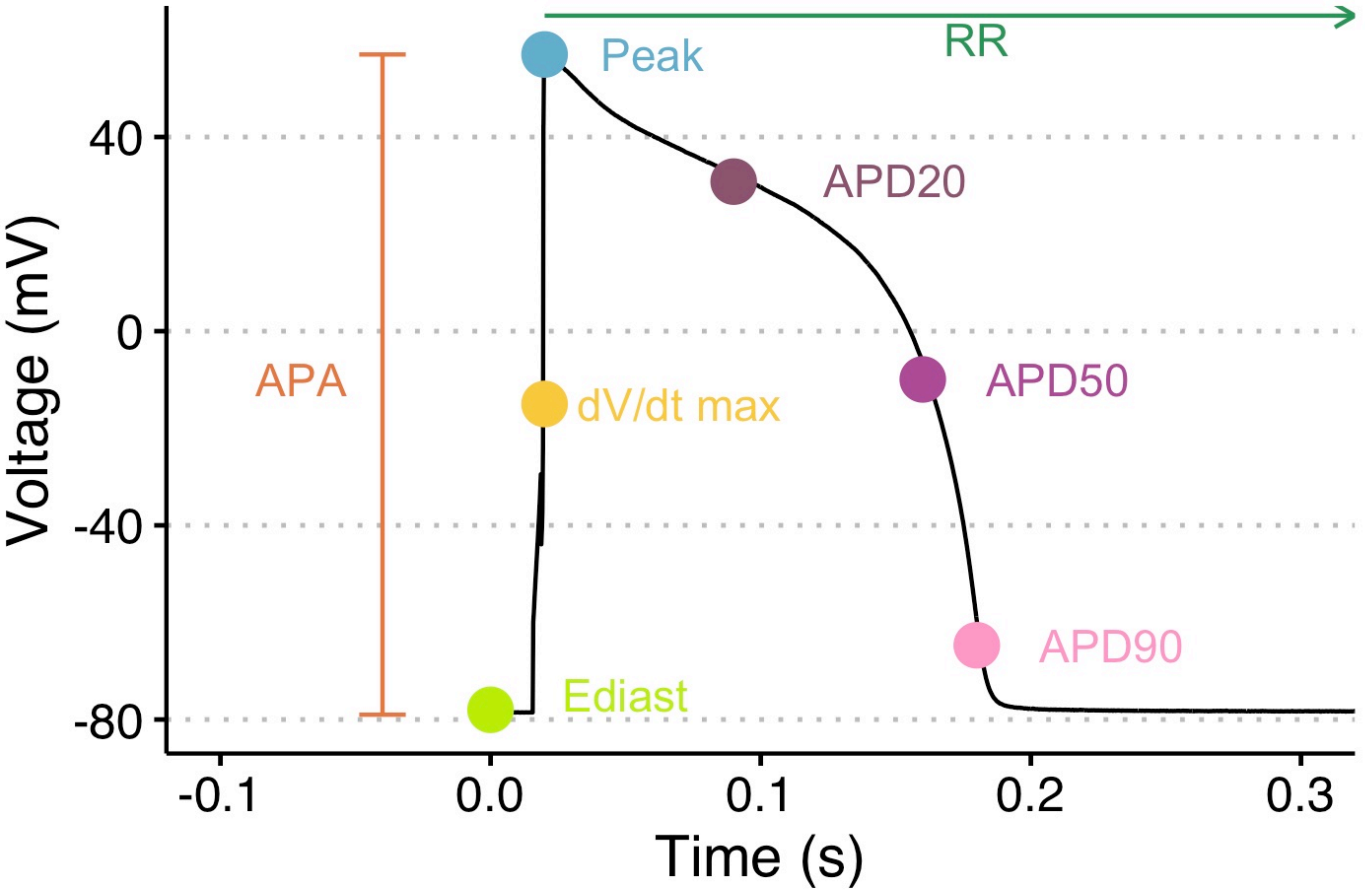
